# Kaposi’s Sarcoma Herpesvirus Is Associated with Osteosarcoma in Xinjiang Uyghur Population

**DOI:** 10.1101/2020.12.11.421719

**Authors:** Qian Chen, Jiangtao Chen, Yuqing Li, Dawei Liu, Yan Zeng, Zheng Tian, Akbar Yunus, Yong Yang, Jie Lu, Xinghua Song, Yan Yuan

## Abstract

Osteosarcoma is the most common malignant tumor of bone predominately affecting adolescents and young adults. Viral etiology of osteosarcoma has been proposed more than a half-century ago but never been proven by identifying any virus authentically associated with human osteosarcoma. The Uyghur ethnic population in Xinjiang China has an unusually high prevalence of Kaposi’s sarcoma-associated herpesvirus (KSHV) infection and elevated incidence of osteosarcoma. In the current study, we explored the possible association of KSHV infection and osteosarcoma occurrence. Our seroepidemiological study revealed that KSHV prevalence was significantly elevated in osteosarcoma patients versus the general population in the Xinjiang Uyghur population (OR, 10.23; 95%CI, 4.25, 18.89). The KSHV DNA genome and viral latent nuclear antigen LANA were detected in most osteosarcoma tumor cells. Gene expression profiling analysis showed that KSHV positive osteosarcoma represents a distinct subtype of osteosarcomas with viral gene-driven signaling pathways that are important for osteosarcoma development. We conclude that KSHV infection is a risk factor for osteosarcoma and KSHV is associated with some osteosarcomas, representing a newly identified viral-associated endemic cancer.

**Significance:** Viral etiology of osteosarcoma was proposed previously but has never been proven by identifying any virus that is authentically associated with human osteosarcoma. The current study revealed an association of human osteosarcoma with KSHV infection in Uyghur osteosarcoma patients. First, this study provides the first evidence that supports the possible viral etiology of human osteosarcoma. The gene expression profiling study showed that KSHV-positive osteosarcoma represents a distinct subtype of osteosarcomas, which is of diagnostic, prognostic and therapeutic significance. Second, KSHV-associated osteosarcomas preferentially occur in children and young adults, predicting that KSHV-positive children in KSHV endemic region may be at great risk for osteosarcoma. Third, the finding extended the range of human cancers associated with viruses.

## Introduction

Osteosarcoma is the most common malignant tumor of bone with an incidence of approximately three cases per million annually worldwide, predominately affecting adolescents and young adults with a major peak between 10 and 14 years old and a second smaller peak in the geriatric population (1, 2). Osteosarcomas have a rather heterogeneous genetic profile and lack any consistent unifying event that leads to its pathogenesis. It may result from oncogenic events sustained by cells in the differentiation lineage hierarchy from mesenchymal stem cells through to osteoblast. Still, it may also suggest that osteosarcoma could arise with multiple etiologies and different pathogenesis. Concerning the etiology of osteosarcoma, three primary potential etiological agents, namely chemical agents, physical agents, and viruses, are believed to associate with osteosarcoma. Numerous chemicals are known to induce osteosarcoma, such as beryllium and methylcholanthrene (3, 4). Radiation exposure is also related to osteosarcoma development (5, 6). The viral etiology for osteosarcoma has been suggested since Finkel isolated viruses from mice that produced similar tumors when injected into newborn mice (7). There was also evidence of a bone tumor virus in the human disease as injection of cell-free extracts of human bone cancer into newborn Syrian hamsters induced various mesenchymal tumors including osteosarcoma (8). However, viral etiology has never been proven by the identification of any virus that is authentically associated with human osteosarcoma. The current study aimed to explore if any human osteosarcoma is associated with a viral infection.

Kaposi’s sarcoma-associated herpesvirus (KSHV), also termed human herpesvirus type 8 (HHV-8), was first identified in Kaposi’s sarcoma (KS) lesions in 1994 (9). KSHV can be found in almost 100% of KS lesions, regardless of their source or clinical subtype (i.e., classic, AIDS-associated, African endemic, or post-transplant KS). Additionally, KSHV is also associated with two lymphoproliferative diseases, namely primary effusion lymphoma (PEL) and multicentric Castleman’s disease (MCD) (10 – 12). We and others showed that human mesenchymal stem cells (MSCs) are highly susceptible to KSHV infection and infection promotes multi-lineage (osteogenic, adipogenic and angiogenic) differentiation (13 – 15). Several lines of evidence suggest that the KSHV-infection of MSCs leads to KS through a mesenchymal-to-endothelial transition (MEndT) process (13, 16).

Given that osteosarcomas originate from mesenchymal stem cells or their immediate lineage progenitors (17, 18) and KSHV can effectively infect MSCs and drive osteogenic differentiation (13), the question was raised if KSHV infection of MSCs contributes to osteosarcoma development. Towards this end, we investigated the association of KSHV infection and osteosarcoma occurrence in the ethnic Uyghur population in Xinjiang Uyghur autonomous region of China, where there is a high seroprevalence of KSHV, a high incidence of KS and a high occurrence of osteosarcoma among the Uyghur population (19 – 21). We found that the seroprevalence of KSHV in Uyghur osteosarcoma patients is significantly higher than that of the Uyghur general population. KSHV genomic DNA and viral latent nuclear antigen (LANA) were detected in osteosarcoma tumors of most of the patients who were KSHV seropositive. Furthermore, gene expression profile analysis of osteosarcoma clinical samples demonstrated that KSHV infection regulates the genes and signaling pathways essential for osteosarcoma development. These results revealed a strong association between KSHV infection and osteosarcoma.

## Results

### Sero-epidemiological evidence for association of KSHV infection with osteosarcoma in Xinjiang Uyghur ethnic population

Xinjiang, China is an endemic area for Kaposi’s sarcoma (KS) and classic KS is prevalent among the ethnic Uyghur population (19). The seroprevalence of KSHV among the Xinjiang Uyghur population is unusually high, ranging from 20.7 to 40.4% compared to that of the general population of China (11.3%) (20). It is striking that although the Uyghurs account for 45% of the total Xinjiang population (according to the 2010 Census of Xinjiang), 68% of osteosarcoma patients diagnosed in our hospital (the First Affiliated Hospital, Xinjiang Medical University) happened to be Uyghurs. It raised a question if KSHV infection is associated with at least some osteosarcomas. To obtain epidemiological evidence regarding this, we compared the KSHV seroprevalence between osteosarcoma patients and the general population. A diagnostic enzyme-linked immunosorbent assay (ELISA) was developed with three recombinant proteins of KSHV, namely LANA, ORF65 and K8.1, and used to examine sera of 21 Uyghur osteosarcoma patients and 327 control individuals (the general Uyghur population). Seventeen of 21 (81%) Uyghur osteosarcoma patients were found seropositive to at least one of the KSHV antigens (*SI Appendix*, Table S1). The serum samples of osteosarcoma patients were also examined in a blinded fashion by an immunofluorescence assay with BCBL-1 cells and LANA nuclear staining with distinctive punctate dots was detected with sera of 17 out of 21 patients, consistent with the results of the ELISA (Fig. 1). In contrast, the control group sera showed a seropositive rate of 29.4% (Table 2; *SI Appendix*, Table S2), consistent with the data previously reported (19, 20). Odds ratio (OR = 10,23, 95%CI: 4.25, 18.89) and P-value (P < 0.0001) indicate that KSHV infection is a risk factor for osteosarcoma occurrence in Xinjiang Uygur population.

**Fig. 1.**
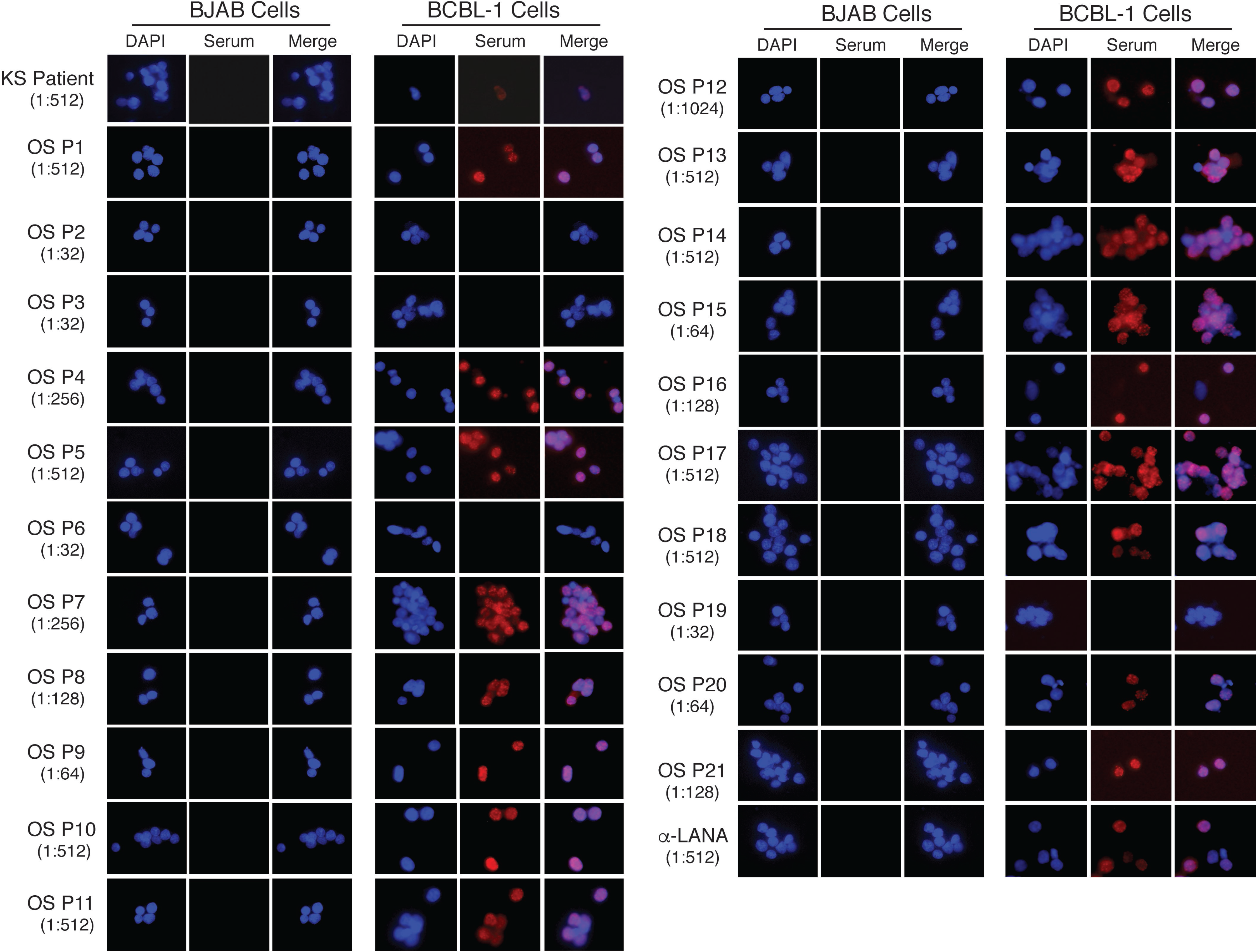
The presence of antibodies specific to KSHV in sera of osteosarcoma patients. Serum samples from 21 Uyghur osteosarcoma patients were examined for KSHV seropositivity with KSHV-negative BJAB and KSHV latently infected BCBL-1 cells. Serum from a Kaposi’s sarcoma patient and an anti-LANA antibody were included as positive controls. Sera and antibody were diluted in a two-fold serial fashion and reacted with the cells on slides (the dilution for each serum was listed under the patient identity number on the left). No serum was found to react with BJAB cells, but some osteosarcoma sera reacted with BCBL-1 cells showing bright punctate staining in the nuclei, a typical LANA staining reported previously (29). The nuclear punctate staining pattern can also be seen in BCBL-1 cells staining with an anti-LANA antibody, which was illustrated in the last row of the figure.

### The KSHV genomic DNA and LANA protein can be detected in most of the osteosarcoma tumors of the KSHV seropositive patients

Surgical specimens of osteosarcoma tumors and adjacent normal tissue samples were obtained from 17 of these 21 patients. These included 14 cases of KSHV seropositive and three seronegative patients. These samples were examined in a blinded fashion for the presence of the KSHV genome in these tumors (the examiners were unaware of patient identities and sample types). Nested PCR was employed to detect KSHV genomic DNA using five sets of primers targeting viral genes K5, ORF25, ORF26, ORF37 and ORF73 (LANA) (Fig. 2A). Quantitative real-time PCR was also performed using ORF73 specific primers in a blinded fashion (Fig. 2B). With 100% consistency, both assays showed that the KSHV DNA genome was detected in 12 out of 14 tumors from the KSHV-seropositive patients, and the other two, namely P14 and P16, had tumors in which KSHV DNA is absent or below the detection of the PCR. The KSHV genome was not detected in the tumors from three KSHV-seronegative patients. No KSHV DNA sequence was detected in adjacent normal tissues of all cases, regardless of their KSHV serological status. Two sets of primers for the EBV genome were included in the Nested PCR assay. No EBV DNA was detected in any of these tumors (Fig. 2A).

**Fig. 2.**
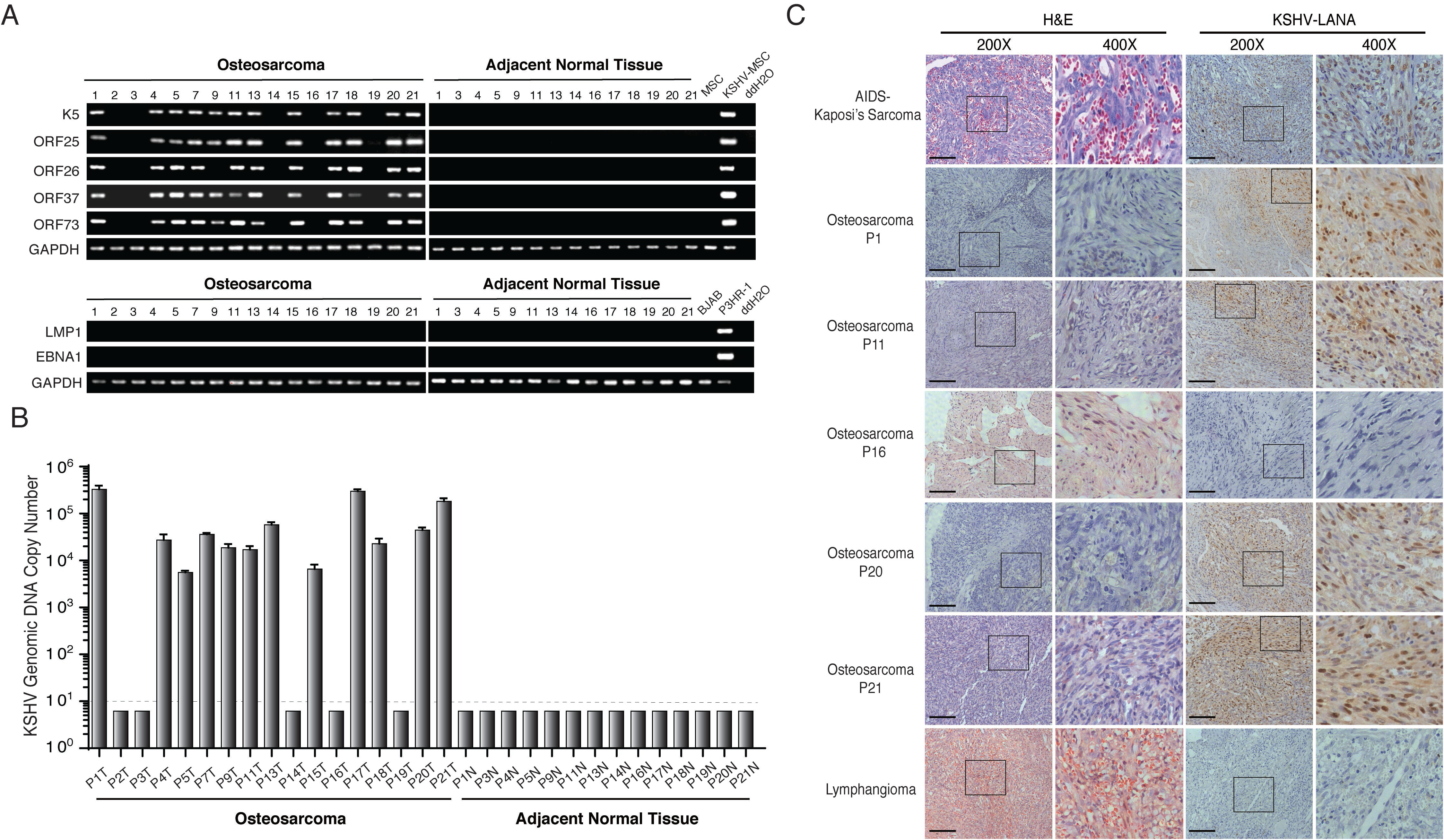
KSHV genomic DNA and latent nuclear antigen (LANA) in osteosarcoma tumors. (A) Detection of KSHV genomic DNA in osteosarcoma tumors using a nested PCR approach. Total DNA was isolated from osteosarcoma tumors and adjacent normal tissues. Two hundred nanogram of each sample was subjected to nested PCR with primers specific to ORFs K5, 25, 26, 37, and 73 (LANA). Primers for EBV LMP1 and EBNA1 are included as a reference. (B) Detection of KSHV genomic DNA in osteosarcoma tumors by quantitative real-time PCR with a pair of primers specific to ORF73 (LANA). The absolute quantification of the KSHV DNA genome (copy number) of each clinical specimen is illustrated. A dashed line shows the limit of detection of the quantitative PCR. (C) Osteosarcoma tumors were subjected to immunohistochemical staining with an antibody against KSHV nuclear antigen LANA. An AIDS-KS tumor sample and a lymphangioma sample were included as controls.

Osteosarcoma samples of five patients were subjected to immunohistochemical analysis for KSHV latent nuclear antigen (LANA). P1, P11, P20 and P21 osteosarcomas exhibited intense LANA staining in spindle- or cigar-shaped osteosarcoma cells (Fig. 2C), indicating that these osteosarcoma cells carry latently infected KSHV. The osteosarcoma of P16 did not express LANA, which was consistent with the fact that the KSHV genome was not detected in the tumor of this patient.

### Gene expression profiling reveals that KSHV-positive osteosarcoma represents a distinct subtype of osteosarcomas

The gene expression profiles of osteosarcoma samples were characterized to reveal possible role of KSHV in osteosarcoma development. Total RNAs were purified from six KSHV-positive (P1, P5, P9, P11, P20 and P21) and four KSHV-negative osteosarcomas (P2, P3, P14 and P16) along with related adjacent normal tissues (except P2 that lacks its adjacent normal tissue) and subjected to RNA-seq analysis. RNA-seq reads were first mapped to the KSHV genome (GQ994935.1) (*SI Appendix*, Table S3) and visualized on a linear scale to provide an overview of highly expressed regions of the genome. At first glance, six KSHV-positive osteosarcomas exhibit two distinct types of viral gene expression pattern: (i) P1, P11 and P20 expressed a high level of PAN RNA; (ii) P5, P9, and P21 expressed a relatively low level of PAN RNA, but a high level of K2 (vIL-6). In addition, the expression of ORF4 (KCP), ORF45 and ORF50 (RTA) were also detected in these tumors (Fig. 3A). Therefore, KSHV-positive osteosarcomas can be grouped into two classes based on the viral gene expression profiles, namely the PAN class and the vIL-6 class. The complete range of read depths across the KSHV genome is visualized on a log scale (*SI Appendix*, Fig. S1).

**Fig. 3.**
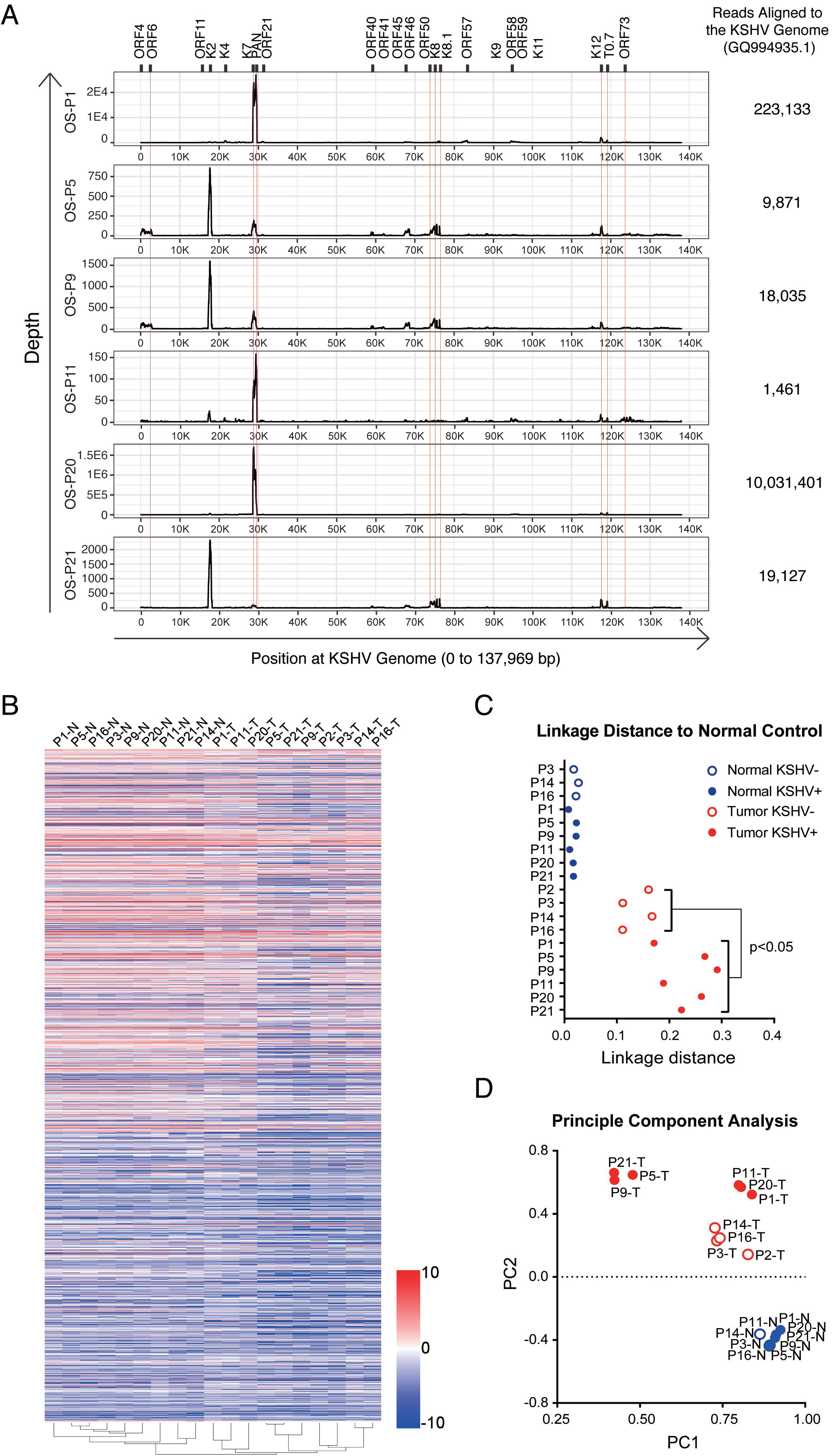
KSHV-positive and -negative osteosarcomas represent distinct subtypes of osteosarcoma on gene expression profiles. (A) KSHV transcriptome of six KSHV-positive osteosarcomas (P1, P5, P9, P11, P20, and P21). RNA-seq reads were aligned to the complete KSHV reference genome (version GQ994935.1) and visualized on a linear scale. The Y-axis represents the number of reads aligned to each nucleotide position of the KSHV genome. KSHV genome positions are indicated at the bottom of each panel. Total reads mapped to the KSHV genome of each clinical sample are listed on the right. (B) The RNA-seq reads of ten osteosarcomas as well as their adjacent normal tissue samples were mapped to the human reference genome (version hg19/GRCh37). Unsupervised clustering of osteosarcoma samples (X-axis) and genes (Y-axis) were performed by the average linkage method. (C) Linkage distance to normal tissue was determined by the Pearson correlation coefficient. (D) The first two principal components of these data were identified and shown in multiple dimensional scaling (MDS) plot.

Then the RNA-seq reads were aligned to the human genome (hg19/GRCh37) and the FPKMs (Fragments Per Kilobase Million) were subjected to unsupervised clustering and differential expression analyses. Clustering of cell samples (X-axis) and genes (Y-axis) were performed by hierarchical clustering with average linkage method and Euclidean distance metric (Fig. 3B). The convergence and divergence among these osteosarcomas and their adjacent normal tissues were determined by linkage distance based on the Pearson correlation coefficient and the principal component analysis (PCA). Results show that the gene expression profiles of all ten osteosarcomas are distinct from the adjacent normal tissues. Four KSHV-negative osteosarcomas are very similar in their gene expression profiles but distantly categorized from KSHV-positive tumors (p<0.05) (Fig. 3C and D), suggesting that KSHV-positive osteosarcomas arose by different pathogenesis than that of non-viral osteosarcomas. Furthermore, KSHV-positive osteosarcomas were divided into two categories visualized by PCA on their cellular gene expression profiles (Fig. 3D). Interestingly, these two categories respectively correspond to the PAN and the vIL-6 classes, suggesting that the different viral gene expression pattern leads to divergent cell reprogramming through either directly affecting host gene expression or changing the environment of host cells.

Differentially expressed genes (DEGs) of osteosarcomas versus their adjacent normal tissues were subjected to a Gene Ontology (GO) analysis to reveal specific and significant associations with specific GO terms. KSHV-positive and -negative osteosarcomas share some common characteristics in gene expression but exhibit more diversity (*SI Appendix*, Fig. S2A). For instance, all categories of osteosarcoma exhibit enriched “multicellular organism development”, “angiogenesis” and “Extracellular matrix organization” Biological Process categories. However, “inflammatory response” and “positive regulation of cell migration” are unique to KSHV-positive osteosarcomas, while “DNA repair” and “response to organic substance” terms are only seen in KSHV-negative osteosarcomas (*SI Appendix*, Fig S2 B – D). Besides, common and unique terms of “Biological Process” or “Molecular Function” are also found between two KSHV-positive classes (PAN-class and vIL6-class) (*SI Appendix*, Fig. S2). It is worth noting that “inflammatory response” is significantly enriched in KSHV-positive osteosarcoma, both PAN-class (2.73% DEGs involved, p = 0.006) and vIL6-class (2.78% DEGs involved, p = 0.0021), but not in KSHV-negative osteosarcomas (*SI Appendix*, Fig S2. B – D). Furthermore, “positive regulation of interferon-gamma production” is uniquely enriched in vIL6-class (0.67% DEGs involved, p = 0.024). These data provide additional evidence of the virus burden in KSHV-positive osteosarcomas and suggest elevated inflammation in KSHV-associated osteosarcoma. Overall, our gene expression profiling analysis showed that KSHV-associated osteosarcoma represents a distinct subtype of osteosarcomas with viral gene-driven signaling pathways that are important for osteosarcoma development. KSHV-positive osteosarcoma is a newly identified viral-associated endemic cancer.

## Discussion

Numerous agents have been implicated in osteosarcoma etiology, including chemical, radiation, and virus (3 – 8, 22). Although osteosarcoma can be induced by a virus in experimental animals (7), whether any human osteosarcoma is caused by virus has been a longstanding and intriguing question but remain elusive. In the current study, we found an association of human osteosarcoma with KSHV in the Xinjiang Uyghur population that has an unusually high prevalence of KSHV infection. First, a serological study for KSHV prevalence in Uyghur osteosarcoma patients versus the general Uyghur population provided epidemiological evidence that KSHV infection is a risk factor for osteosarcoma. Second, the KSHV genome and viral latent protein LANA were detected in most osteosarcoma tumors of KSHV-positive patients. Third, gene expression profiling analysis showed that KSHV-positive osteosarcoma represents a distinct subtype of osteosarcoma. Taken together, our results demonstrated that KSHV-positive osteosarcoma is a newly identified viral-associated endemic cancer, presenting the first evidence that human osteosarcoma is associated with an oncogenic virus.

Another important phenomenon revealed in this study is that KSHV-associated osteosarcomas preferentially occur in children and young adults. 86% osteosarcomas of the patients aged under 30 years are KSHV-associated, in contrast to that none of the patients above 30 was found to have KSHV-positive osteosarcoma (P-value = 0.024, Table 1). It has been reported that KSHV infection occurs in early childhood in the Xinjiang Uyghur population (23). Thus, KSHV-positive children may be at significant risk for osteosarcoma, which appears to be true in the Uyghur population. Interestingly, two older adult patients (P14 and P16, aged 53 and 39, respectively) had KSHV-negative osteosarcomas despite being KSHV seropositive. This observation suggests that KSHV-positive osteosarcoma is mainly associated with pediatric patients. In adults, osteosarcoma may, therefore, arise through non-viral etiology even in KSHV-seropositive individuals.

**Table 1.** Baseline Characteristics of Uyghur Osteosarcoma Patients.

| Characteristics | Patient KSHV Sero-status (N = 21) |  |  | Patient KSHV Genomic DNA (N = 17) |  |  |
| --- | --- | --- | --- | --- | --- | --- |
|  | Number of Subjects (%) | KSHV Sero-positivity (%) | P Value | Number of Subjects (%) | KSHV DNA-positivity (%) | P Value |
| Gender |  |  |  |  |  |  |
| Male | 9 (42.9) | 9 (100.0) | 0.104 | 8 (47.1) | 6(75.0) | 1.00 |
| Female | 12 (57.1) | 8 (66.7) |  | 9 (52.9) | 6 (66.7) |  |
| Age group |  |  |  |  |  |  |
| 5-30 years | 18 (85.7) | 15 (83.3) | 0.489 | 14 (82.4) | 12 (85.7) | 0.024 |
| >30 years | 3 (14.3) | 2 (66.7) |  | 3 (17.6) | 0 (0) |  |
| Diagnosis |  |  |  |  |  |  |
| CIMOS <sup>a</sup>, Fibroblastic | 3 (14.2) | 2 (66.6) | 0.962 | 2 (11.8) | 1 (50.0) | 0.478 |
| CIMOS, Osteoblastic | 6 (28.6) | 5 (83.3) |  | 4 (23.6) | 4 (100.0) |  |
| CIMOS, Chondroblastic | 2 (9.5) | 2 (100.0) |  | 2 (11.8) | 2 (100.0) |  |
| CIMOS, Myxoid | 1 (4.8) | 1 (100.0) |  | 0 (0) | 0 (0) |  |
| CIMOS, Round Cell | 1 (4.8) | 1 (100.0) |  | 1 (5.9) | 1 (100.0) |  |
| CIMOS, Mixed | 5 (23.8) | 3 (60.0) |  | 5 (29.4) | 2 (40.0) |  |
| CIMOS, NOS <sup>b</sup> | 1 (4.8) | 1 (100.0) |  | 1 (5.9) | 1 (100.0) |  |
| WDIOS <sup>c</sup> | 2 (9.5) | 2 (100.0) |  | 2 (11.8) | 1 (50.0) |  |
| Tumor location |  |  |  |  |  |  |
| Femur | 11 (52.3) | 9 (81.8) | 0.421 | 8 (47.0) | 4 (50.0) | 0.109 |
| Tibia | 5 (23.8) | 5 (100.0) |  | 5 (29.4) | 5 (100.0) |  |
| Humerus | 2 (9.5) | 2 (100.0) |  | 2 (11.8) | 2 (100.0) |  |
| Pelvis | 1 (4.8) | 0 (0) |  | 1 (5.9) | 0 (0) |  |
| Clacivle | 1 (4.8) | 1 (100.0) |  | 1 (5.9) | 1 (100.0) |  |
| Scapula | 1 (4.8) | 1 (100.0) |  | 0 (0) | 0 (0) |  |
| Chemotherapy |  |  |  |  |  |  |
| Neo-adjuvant Chemotherapy | 15 (71.4) | 12 (80.0) | 1.00 | 12 (70.6) | 8 (66.7) | 1.00 |
| Never | 6 (28.6) | 5 (83.3) |  | 5 (29.4) | 4 (80.0) |  |
| Total | 21 (100.0) | 17 (81.0) |  | 17 (100.0) | 12 (70.6) |  |

<sup>a</sup> CIMOS: Conventional intramedullary osteosarcoma; <sup>b</sup> NOS: No otherwise specified; <sup>c</sup> WDIOS: Well-differentiated intramedullary osteosarcoma.

**Table 2.** Risk of KSHV Infection with Osteosarcoma Occurrence.

|  | Osteosarcoma (%)<br>(N = 21) | Control Group (%)<sup>a</sup><br>(N = 327) | Odds Ratio<br>(95% CI) | P Value |
| --- | --- | --- | --- | --- |
| KSHV-positive | 17 (81.0) | 96 (29.4) | 10.23 (4.25-18.89) | < 0.0001 |
| KSHV-negative | 4 (19.0) | 231 (70.6) |  |  |
| Total | 21 (100.0) | 327 (100.0) |  |  |

<sup>a</sup> Control group includes 327 Uyghur donors who came to the hospital for annual physical examination. KSHV prevalence in Uyghur osteosarcoma patients = 81.0% KSHV prevalence in Uyghur general population = 29.4%

Osteosarcoma is not a single disease but a collection of neoplasms with different etiologies, sharing a histological hallmark of osseous matrix production in association with malignant cells. Our study showed that KSHV-positive and -negative osteosarcomas exhibit distinct gene expression profiles, implicating that osteosarcomas caused by biological (virus), chemical (carcinogens), or physical (radiation) agents can be categorized into different subtypes based on their gene expression profiles and specific GO enrichment. Furthermore, KSHV-positive osteosarcomas can be classified into two distinct classes (the PAN class and the vIL-6 class) based on their viral and cellular gene expression profiles. Further studies are warranted to determine if these gene signature-defined categories are linked to certain clinical features and osteosarcoma manifestation, which will be of great diagnostic or prognostic significance. On the other hand, it is also important to identify gene signatures for pan-osteosarcomas. For example, TGF-β1 and 3, as well as their receptor TGF-βRII, were found to be up-regulated in all osteosarcomas regardless of KSHV infection status. It is consistent with previous observations that high-grade osteosarcomas have a significantly higher expression of TGF-β1, which may influence the aggressive clinical behavior of the sarcoma (24), and that TGF-β3 is associated with disease progression (25). In addition, “multicellular organism development” and “extracellular matrix (ECM) organization” are also found at the top of the significant GO term list for all osteosarcoma samples and present a common feature for all osteosarcomas.

There is geographic variation in the prevalence of KSHV and associated diseases (26). The prevalence of KSHV-associated osteosarcoma, as well as the ratio of osteosarcomas with different etiologies, may vary in different regions upon the causative virus prevalence and ethnic background. In North America and the Han population of China (the ethnic majority of China) where the KSHV prevalence is low, KSHV-associated osteosarcomas are believed to be rare. We collected nine osteosarcoma patients of the Han, and only one was KSHV-seropositive (11%) (*SI Appendix*, Table S4). We analyzed this Han KSHV-positive osteosarcoma (HP2) along with a KSHV-negative tumor (HP9) for their gene expression profiles. Result showed that HP2 osteosarcoma is a PAN-class tumor and the biology of the HP2 osteosarcoma is similar to Uyghur tumors of the same category (P1, P11 and P20), revealed RNA-seq principle component analysis (PCA) (*SI Appendix*, Fig. S3). Although the sample size for osteosarcomas of Han patients is too small to draw any conclusion, the trend suggests that (i) KSHV-associated osteosarcoma may not be the majority of this cancer in the Han population; (ii) once being infected by KSHV at an early age, children of other races can also develop KSHV-associated osteosarcoma. KSHV prevalence is known to be high in specific areas such as Eastern African (Uganda, Cameroon, Democratic Republic of Congo, Tanzania and Zambia). It is very intriguing to know if the occurrence of childhood osteosarcoma is high in this area. Since many developing countries do not have cancer registries or many childhood cancers are not diagnosed, there is a lack of epidemiology study for childhood osteosarcoma in these countries. However, a simulation-based study of global childhood cancer suggested that pediatric osteosarcoma in Eastern Africa is several-fold higher than in Southern and Northern Africa in parallel with the incidence of Kaposi’s sarcoma (27). On the other hand, the possibility also exists that KSHV-associated osteosarcoma is unique in the Xinjiang Uyghur population, analogous to EBV-associated Burkitt’s lymphoma, which is common in African children, and EBV-caused nasopharyngeal carcinoma, which mainly occurs in Southern China. Further investigation is warranted to identify the risk factors associated with the area or population in hope that strategies can be developed to reduce osteosarcoma occurrence such as preventing children from KSHV infection at an early age.

## Materials and Methods

### Patients and participants

The study included patients with osteosarcoma (age from 5 to 53 years-old, median 15 years-old) who were admitted and treated in the period of 2016 – 2019 in the Division of Orthopedic Oncology, the First Affiliated Hospital of Xinjiang Medical University, Urumqi, China. Among the patients, the majority is of Uyghur ethnicity (21 Uyghurs, 1 Kazakh, 9 Hans). All patients underwent a core needle biopsy or surgical biopsy for diagnosis. Blood samples were also collected. Fifteen Uyghur osteosarcoma patients received 4 – 10 weeks neo-adjuvant chemotherapy (Doxorubicin, Cisplatin and Ifosfamide) and six were not treated with chemotherapy prior to surgery. Then extensive resection or radical surgery was performed. Surgically removed osteosarcoma tumors and adjacent normal tissues were collected from these patients. Demographic and clinical data were listed in Table 1 and *SI Appendix* Table S1. Sera were also collected from 327 healthy Uyghur donors who came to the hospital for annual physical examination as reference (*SI Appendix,* Table S2). The human sample collection and the use of clinical samples in the research were approved by the Institutional Review Boards of the Xinjiang Medical University, the First Affiliated Hospital (Approval No. 2018-112903) and Sun Yet-sen University (Approval No. 2015-028). Informed consent was obtained from all osteosarcoma patients (or their guardians) and healthy donors.

### Reagents and antibodies

Cell culture medium (MEM-alpha), streptomycin, penicillin, TRIzol reagent were purchased from Invitrogen. Nonessential amino acids, glutamine, β-glycerophosphate, dexamethasone, Alizarin Red S, paraformaldehyde were obtained from Sigma-Aldrich. Antibody against LANA (ab4103) antibody was purchased from Abcam. Alexa Fluor 555 goat anti-Human IgG (H+L) antibody (A21433) was purchased from Invitrogen. Alexa Fluor 555 goat anti-Rat IgG antibody (A21434) was purchased from Life Technologies. Hematoxylin (G1004) and eosin (G1002) were obtained from Servicebio technology. HRP labeled goat anti-rabbit IgG (SP-D2) and DAB reaction kit (DAB-1031) were purchased from Maxim Biotechnologies.

### Expression and purification of KSHV K8.1, ORF65, LANA proteins in E. coli

cDNAs of KSHV K8.1, ORF65 and LANA were cloned into the pET-28a vector with a hexahistidine (6xHis) tagged at the N-terminus. *E.coli* Rosetta cells were transformed with each of the recombinant plasmids and cultured in LB medium contain 30μg/mL Kanamycin. The expression of these proteins was induced with 1mM isopropyl β-D-thiogalactoside (IPTG) when the optical density of culture reached OD of 0.6. For K8.1, induced culture was collected after 4 hours cultivation at 16°C, while for LANA and ORF65, cultures were collected after 4 hours cultivation at 37°C. Cells were ultrasonicated in lysis buffer containing PMSF without DTT and proteins were purified with Ni-NTA column chromatography. Protein concentrations were determined by BCA protein assay kit (Thermo Fisher).

### Enzyme-Linked Immunosorbent Assay (ELISA)

Purified K8.1, ORF65, and LANA proteins (100 μl, 5 μg/mL) were respectively coated on ELISA plates (Jet Biofil) in coating buffer (0.1 M NaHCO_3_, pH9.6 –10.0) at 4°C overnight. Plates were saturated with blocking buffer (5% dried skimmed milk in PBS containing 0.1% Tween 20). Each serum in a series of dilutions (1:50 to 1:1600) reacted with coated plates at 37°C for 90 min. After washing with PBST three times, a peroxidase-conjugated anti-human IgG antibody (1:3000 dilution) was added and incubated at 37°C for 30min. Tetramethyl-benzidine (TMB) and hydrogen peroxide substrates were dispensed and incubated in the dark at 37°C for 15 min. The plates were read at 450 nm (OD450) using a microplate reader (BioTek) with the cut-off for seropositivity of OD450>0.5. Before this ELISA system was used to analyze osteosarcoma patient sera, the system had been verified with 21 Uyghur sera with known KSHV serological status (LANA, ORF65m K8.1 seropositivity) (28) with 98.4% consistency and accuracy. Each osteosarcoma patient serum was tested three times in a blinded fashion. An ELISA titer of >1:100 was considered to be positive. To ensure inter-assay comparability, we used a “highly positive” serum from the previous assay as a positive control and a negative serum from a healthy donor as a negative control.

### Immunofluorescence Assay (IFA)

BCBL-1 (KSHV latently infected) and BJAB (KSHV-negative) cells were fixed with 3.6% formaldehyde in PBS and permeabilized with 0.1% Triton X-100. Cells were pre-incubated with 1% BSA and then reacted with patient sera in two-fold serial dilutions. AIDS-KS patient serum and an anti-LANA antibody (ab4103, Abcam) were included as positive controls. Alexa Fluor555 goat anti-human IgG (1:500 dilution) was used as the secondary antibody (Alexa Fluor555 goat anti-Rat IgG for anti-LANA control). Cells were visualized under a Zeiss Observer.Z1 fluorescence microscope. The assay was done in a blinded manner, and the serum yielding the nuclear punctate pattern in > 1:64 dilution was considered positive.

### Polymerase chain reaction (PCR) analyses

Total DNA was extracted from each tumor or adjacent normal tissue sample using a HiPure Tissue DNA Mini Kit (Magen). Two hundred nanogram of each DNA was subjected to nested polymerase chain reaction (PCR). The oligonucleotide primers for the first and second-round PCR for KSHV K5, ORF25, ORF26, ORF37, ORF73 (LANA), and EBV LMP1, EBNA1 were listed in *SI Appendix* Table S5. PCR was performed in a 50 μl-volume reaction [0.4 μM primers and 2xPrimeSTAR HS (Premix), 5% DMSO] as follows: denaturing at 95°C for 5 min, 25 cycles of reaction (95°C for 30 sec, 55–62°C for 30 sec, 72°C for 30 sec) and final elongation of 72°C for 7 min. PCR was carried out in a blinded fashion (the examiner was unaware of patient identities and sample types). Each sample was tested three times independently.

### Real-time polymerase chain reaction (PCR) and quantitative RT-PCR analyses

Two hundred nanogram purified tissue DNA was subjected to quantitative real-time PCR with KSHV ORF73 primers and SYBR Green Master Mix (ThermoFisher) on LightCycler 480II (Roche). LANA standard DNA in serial dilutions were analyzed simultaneously with osteosarcoma DNAs for a standard curve. KSHV genomic DNA copy numbers of initial specimens were calculated according to the standard curve. The lowest standard DNA copy number used to construct the standard curve (10 DNA copies) still exhibited positive peak and was in the linear range in the standard curve, 10 copies is set to be the sensitive line for our assay. Ct value cutoff is 35 cycles. For the quantification of RNA, total RNA was isolated using TRIzol reagent (Invitrogen). RNA concentration was measured by Nanodrop 2000 (ThermoFisher). Five hundred nanogram RNA was reverse-transcribed into cDNA followed by real-time PCR with SYBR Green Master Mix and specific primers on LightCycler 480II. The primer sequences used are listed in *SI Appendix* Table S5.

### Immunohistochemical (IHC) analysis

Osteosarcoma clinical samples were fixed in 4% Paraformaldehyde (PFA) and decalcified in 10% EDTA/0.2% PFA in PBS in microwave decalcifying apparatus. Complete decalcification was verified by X-ray images. Tumor samples, including Osteosarcoma, Kaposi’s sarcoma and Lymphangioma, were impregnated in paraffin and sections were subjected to hematoxylin and eosin (H&E) and immunohistochemical (IHC) staining. For IHC, after removal of endogenous peroxidase with 3% H_2_O_2_ and rinsing in PBS, sections were incubated with an antibody against LANA (ab4103, Abcam) in 1:100 dilution at 4°C overnight. A goat anti-rabbit HRP secondary antibody (DAB-1031, Maxim) was used, followed by metal enhanced DAB colorimetric detection. Then sections were counterstained with hematoxylin.

### RNA sequencing (RNA-seq)

Osteosarcoma tumors and adjacent normal tissues were lysed and total RNA was extracted with TRIzol reagent. DNA contamination was eliminated by DNase I treatment. RNA purity was determined using the Qubit 3.0 Fluorometer (Life Technologies). For each sample, 1-3 μg RNA was used to generate sequencing libraries with NEBNext Ultra™ RNA Library Prep Kit for Illumina (#E7530L, NEB, USA) with poly(A) Magnetic Isolation Module (NEB#7490). Fragmentation was carried out using divalent cations under elevated temperature in NEBNext first strand synthesis reaction buffer. First-strand cDNA was synthesized using random hexamer primer and RNase H. Second strand cDNA was synthesized using DNA polymerase I and RNase H. The library fragments were purified with QiaQuick PCR kits. Index codes were added to attribute sequences to each sample. The libraries were sequenced on an Illumina Hiseq 4000 platform and 150 bp paired-end reads were generated.

### Data process

Raw data in fastq format were filtered, mapped and analyzed as before.13 Briefly, the short reads were aligned to the KSHV reference genome (version GQ994935.1) and the human reference genome (version hg19/GRCh37). The quality of raw data was viewed by multiQC. The number of clean tags mapped to each gene was counted by FPKM (Fragments per Kilo bases per Million reads). Corrected p-value (q-value) < 0.05 and |log2 (fold change) | > 1 were set as threshold for significantly different expression.

### Data analysis

Gene Ontology (GO) analysis was performed with GO tools (http://www.geneontology.org/), and GO terms with p-value < 0.05 were considered as significantly enriched gene sets. Venn diagrams were drawn by Venny 2.0 online (http://bioinfogp.cnb.csic.es/tools/venny/index.html). Unsupervised clustering of samples (X-axis) was performed with the average linkage method and Euclidean distance metric, respectively. Linkage distance was performed by calculating the Pearson correlation coefficient with Normal control and then subtracted by 1. GSEA was performed using GSEA-3.0 software between osteosarcoma tumor and adjacent normal tissue samples with 2000 of “Geneset” permutations type and default values for other parameters. FPKM and RPKM values of RNA expression were used in this analysis. KEGG, Reactome, and hallmark gene sets were used in this analysis (http://software.broadinstitute.org/gsea/msigdb/collections.jsp). All differentially expressed pathways with FDR q-value < 0.1 were kept for subsequent analysis.

### Statistical analyses

Data were analyzed with SPSS 25.0 program. Fisher’s exact test was used to analyze baseline data. Odds Ratio (OR) was calculated to estimate the association between KSHV infection and osteosarcoma occurrence. Woolf’s method was applied to calculate a 95% confidence interval. P values <0.05 were considered significant (*P<0.05, **P<0.01, ***P<0.001).

## Data availability

RNA-seq data obtained in this study are available in NCBI GEO (Gene Expression Omnibus) database, accession: GSE126209.

## Acknowledgments

We thank Yuan Lab members for discussion, constructive suggestions, and participation in blinded serological and quantitative PCR experiments. The research reported in this publication was supported by the Natural Science Foundation of China (81530069).

## Supplementary Information

**This PDF file includes**

Table S1. Demographic Characterization and KSHV Positivity of 21 Uyghur Osteosarcoma Patients.

Table S2. Baseline Characteristics of 327 Control Participants.

Table S3. Osteosarcoma RNA-seq Raw Reads and Alignment to the KSHV and Human Genomes.

Table S4. Demographic Characterization and KSHV Positivity of 9 Han Osteosarcoma Patients.

Table S5. Oligonucleotides Used in the Study.

Figure S1. KSHV Transcriptome of Six KSHV-Positive Osteosarcoma in Log Scale.

Figure S2. Gene Ontology (GO) Analysis of KSHV-positive and KSHV-negative Osteosarcomas.

Figure S3. Comparison of Han Osteosarcoma with Six Uyghur Osteosarcomas on KSHV Transcription Profile.

Figure S4. KSHV-positive and -negative Osteosarcomas, Regardless of Han and Uyghur Patients, represent distinct subtypes of osteosarcoma on Gene Expression Profile.

**Supporting Information – Table S1.** Demographic Characterization and KSHV Positivity of 21 Uyghur Osteosarcoma Patients.

| Patient Code | Sex | Age | Diagnosis | Tumor Location | Ethnicity | Serology ELISA |  |  | IFA KSHV | KSHV Genome in Tumor |
| --- | --- | --- | --- | --- | --- | --- | --- | --- | --- | --- |
|  |  |  |  |  |  | K8.1 | ORF65 | LANA |  |  |
| P1 | M | 28 | CIMOS, NOS | Left humerus | Uyghur | - | + | + | + | + |
| P2 | F | 15 | CIMOS, Mixed | Left pelvis | Uyghur | - | - | - | - | - |
| P3 | F | 46 | CIMOS, Fibroblastic | Left femur | Uyghur | - | - | - | - | - |
| P4 | F | 8 | CIMOS, Osteoblastic | Right tibia | Uyghur | + | - | + | + | + |
| P5 | M | 22 | CIMOS, Chondroblastic | Right tibia | Uyghur | + | + | + | + | + |
| P6 | F | 17 | CIMOS, Osteoblastic | Right scapula | Uyghur | - | - | - | - | N.S. |
| P7 | F | 30 | CIMOS, Mixed | Right clavicle | Uyghur | - | + | + | + | + |
| P8 | F | 27 | CIMOS, Osteoblastic | Left femur | Uyghur | - | - | + | + | N.S. |
| P9 | M | 10 | CIMOS, Osteoblastic | Left femur | Uyghur | - | - | + | + | + |
| P10 | M | 20 | CIMOS, Fibroblastic | Right femur | Uyghur | - | + | + | + | N.S. |
| P11 | F | 10 | CIMOS, Chondroblastic | Left tibia | Uyghur | - | - | + | + | + |
| P12 | F | 13 | CIMOS, Myxoid | Left femur | Uyghur | + | + | + | + | N.S. |
| **P13** | F | 11 | CIMOS,<br>Round cell | Left femur | Uyghur | + | + | + | + | + |
| **P14** | M | 53 | WDIOS | Left femur | Uyghur | + | + | + | + | - |
| **P15** | F | 28 | WDIOS | Left femur | Uyghur | - | - | + | + | + |
| **P16** | M | 39 | CIMOS, Mixed | Left femur | Uyghur | - | + | + | + | - |
| **P17** | F | 9 | CIMOS,<br>Osteoblastic | Right tibia | Uyghur | + | + | + | + | + |
| **P18** | M | 9 | CIMOS,<br>Fibroblastic | Left humerus | Uyghur | + | + | + | + | + |
| **P19** | F | 14 | CIMOS, Mixed | Right femur | Uyghur | - | - | - | - | - |
| **P20** | M | 15 | CIMOS,<br>Osteoblastic | Left femur | Uyghur | - | - | + | + | + |
| **P21** | M | 15 | CIMOS, Mixed | Left tibia | Uyghur | + | - | + | + | + |

CIMOS: Conventional intramedullary osteosarcoma; WDIOS: Well-differentiated intramedullary osteosarcoma; NOS: No otherwise specified; +: Positive; - : Negative; F: Female; M : Male; N.S. : No Sample.

**Supporting Information – Table S2.** Baseline Characteristics of 327 Control Participants (General Uyghur Population)

| Characteristics | Number of subjects (%) | KSHV sero-positivity (%) | P Value |
| --- | --- | --- | --- |
| Gender |  |  |  |
| Male | 142 (43.4) | 46 (32.4) | 0.291 |
| Female | 185 (56.6) | 50 (27.0) |  |
| Age group |  |  |  |
| 5-30 years | 114 (34.9) | 28 (24.6) | 0.164 |
| >30 years | 213 (65.1) | 68 (31.9) |  |
| Smoking tobacco |  |  |  |
| Never | 229 (70.0) | 65 (28.4) | 0.555 |
| Ever | 98 (30.0) | 31 (31.6) |  |
| Drinking Alcohol |  |  |  |
| No | 189 (57.8) | 53 (28.0) | 0.541 |
| Yes | 138 (42.2) | 43 (31.2) |  |
| Total | 327 (100.0) | 96 (29.4) |  |

Age (Years) Mean ± SD): 38.37 ± 13.40; Minimum – maximum: 9 – 86.

**Supporting Information – Table S3.** Osteosarcoma RNA-seq Raw Reads and Alignment to the KSHV and Human Genomes.

Age (Years) Mean ± SD): 38.37 ± 13.40; Minimum – maximum: 9 – 86. Supporting Information – Table S3. Osteosarcoma RNA-seq Raw Reads and Alignment to the KSHV and Human Genomes.
| Sample | Patient ID | Total Reads | Aligned to KSHV Transcriptome | Aligned to Human Transcriptome |
| --- | --- | --- | --- | --- |
| **Osteosarcoma** | P1 | 63,953,856 | 223,133 | 60,491,393 |
|  | P2 | 69,061,060 | 1 | 65,452,961 |
|  | P3 | 81,498,617 | 8 | 79,602,603 |
|  | P5 | 72,040,573 | 9,871 | 68,864,981 |
|  | P9 | 70,845,288 | 18,035 | 66,470,038 |
|  | P11 | 60,649,859 | 1,461 | 57,958,356 |
|  | P14 | 75,958,617 | 3 | 74,449,359 |
|  | P16 | 75,675,293 | 5 | 74,067,075 |
|  | P20 | 97,844,318 | 10,031,401 | 83,058,763 |
|  | P21 | 98,122,848 | 19,127 | 94,631,752 |
|  | HP2 | 43,313,298 | 1,646,210 | 39,515,009 |
|  | HP9 | 45,779,360 | 234 | 42,938,247 |
| **Adjacent Normal Tissue** | P1 | 65,287,890 | 2,717 | 62,538,711 |
|  | P3 | 77,402,745 | 3 | 75,688,331 |
|  | P5 | 79,508,710 | 5 | 77,772,027 |
|  | P9 | 72,239,269 | 7 | 70,439,146 |
|  | P11 | 59,988,497 | 40 | 57,712,755 |
|  | P14 | 77,691,776 | 66 | 76,247,965 |
|  | P16 | 95,230,519 | 2 | 93,522,293 |
|  | P20 | 86,440,986 | 28 | 84,727,078 |
|  | P21 | 96,247,969 | 112 | 94,370,220 |
|  | HP2 | 47,348,598 | 2 | 45,446,160 |
|  | HP9 | 37,135,030 | 6 | 35,434,755 |

**Supporting Information – Table S4.**
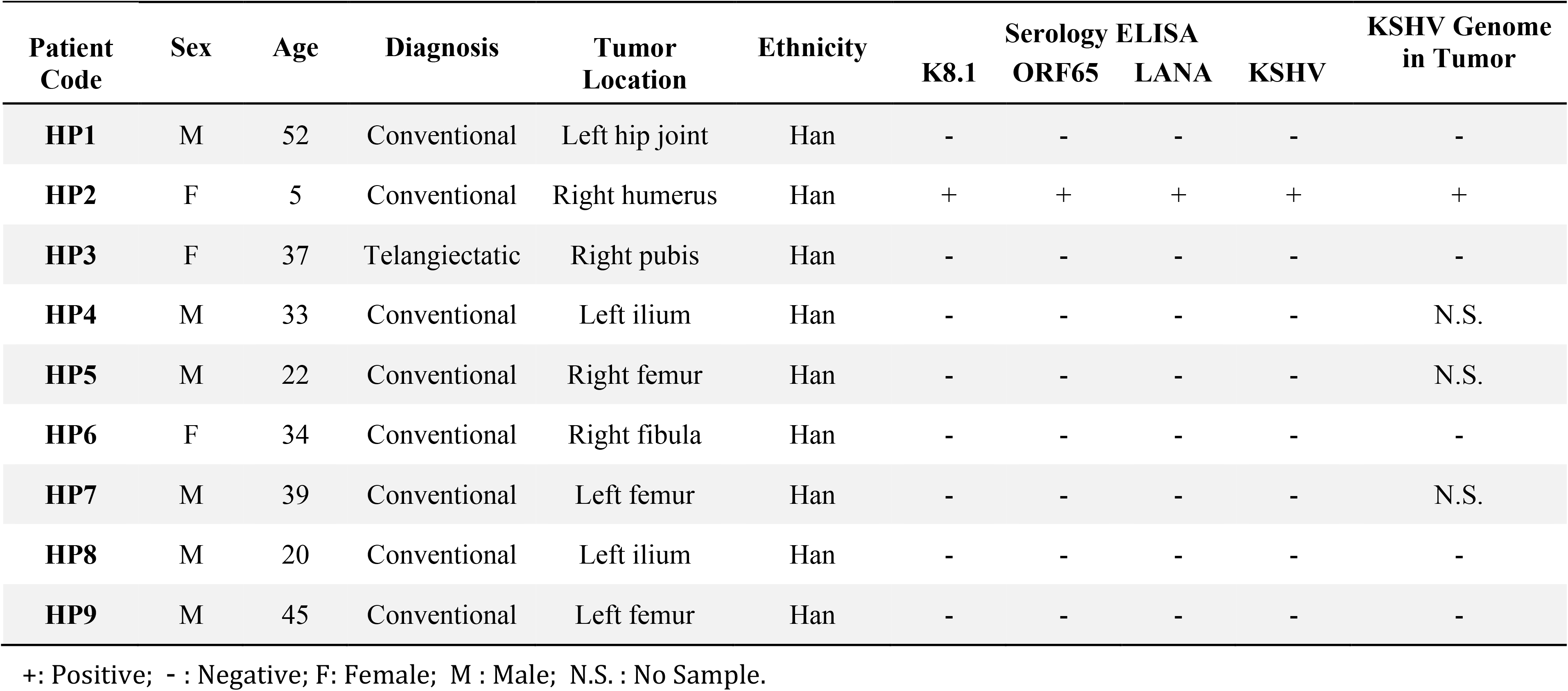
Demographic Characterization and KSHV Positivity of 9 Han Osteosarcoma Patients.

**Supporting Information – Table S5.**
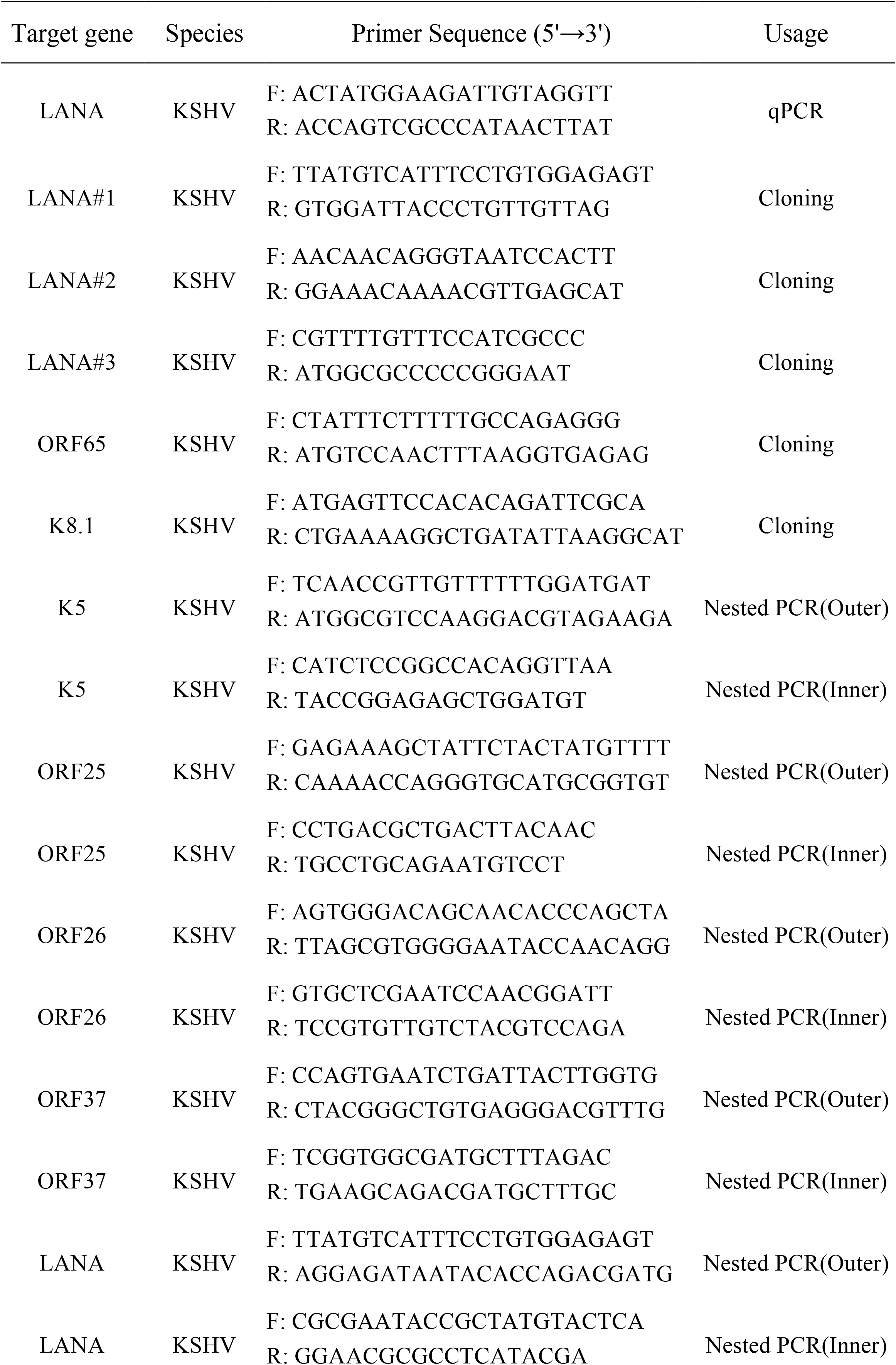
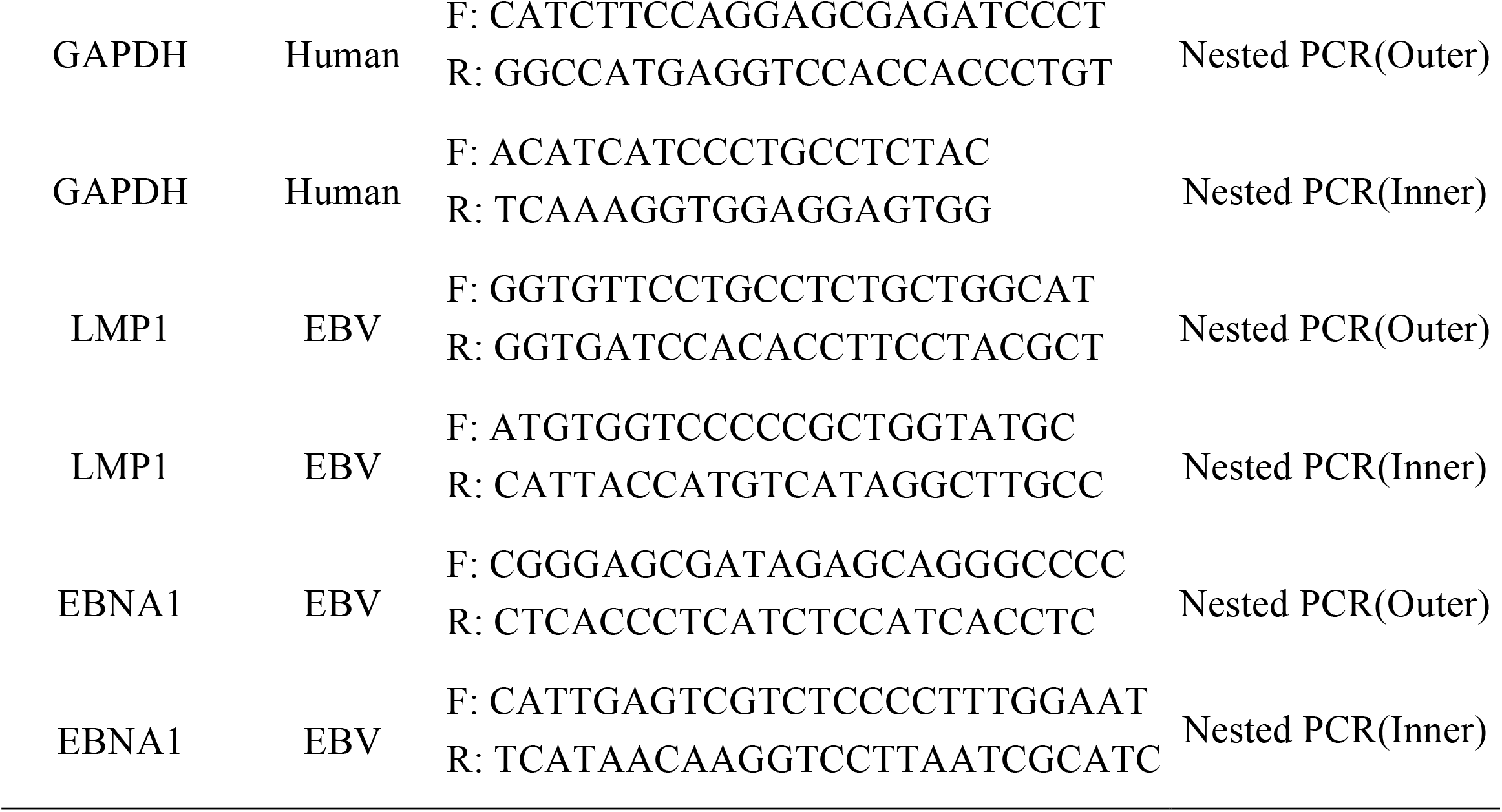
Oligonucleotides used in the study.

**Supporting Information – Fig. S1.**
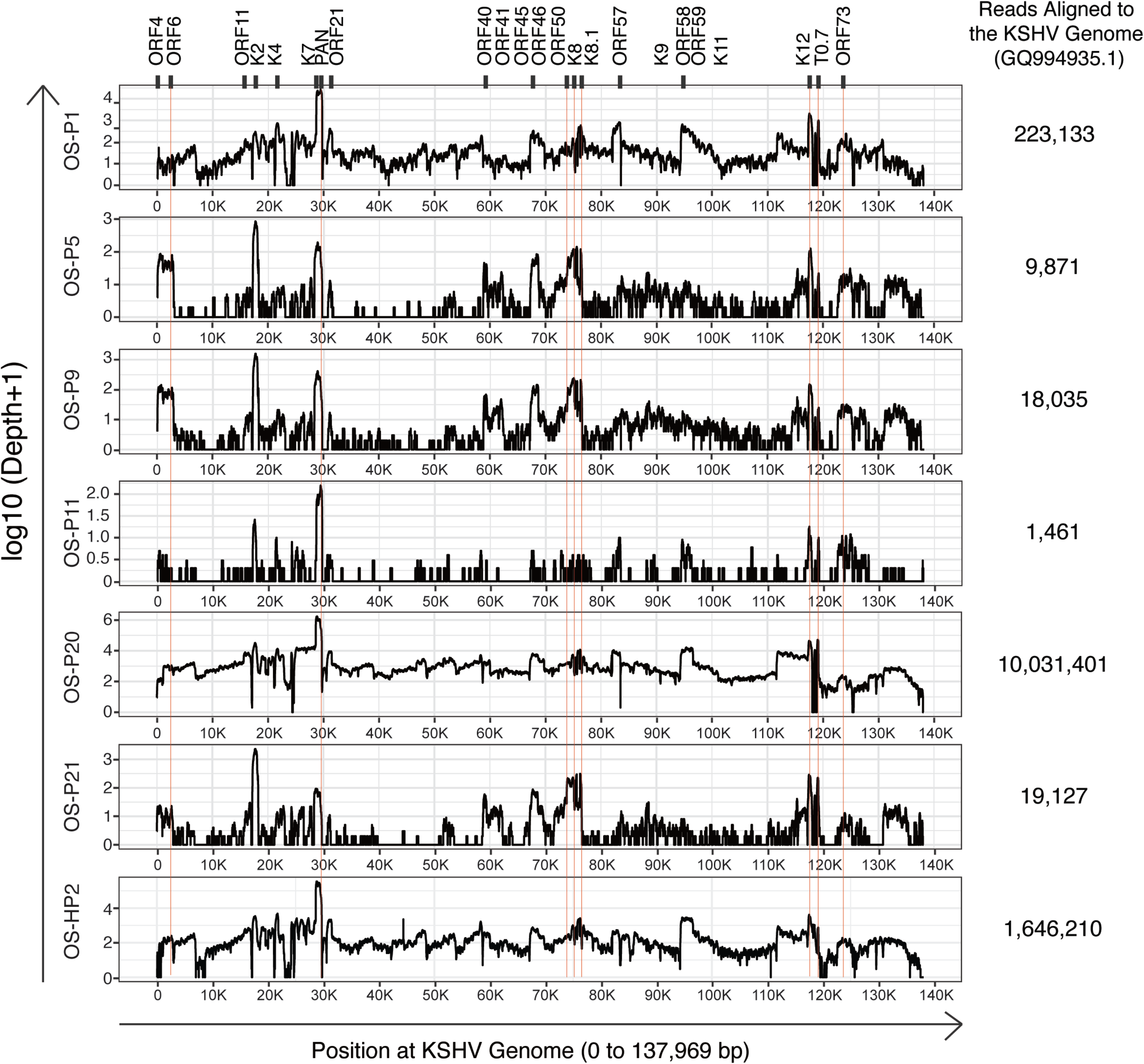
KSHV transcriptome of six KSHV-positive osteosarcomas on logarithmic scale. RNA-seq reads were aligned to the complete KSHV reference genome (version GQ994935.l). The Y-axis represents the number of reads aligned to each nucleotide position ofthe KSHV genome in a log-10 scale. Total reads mapped to the KSHV genome of each clinical sample are listed on the right.

**Supporting Information – Fig. S2.**
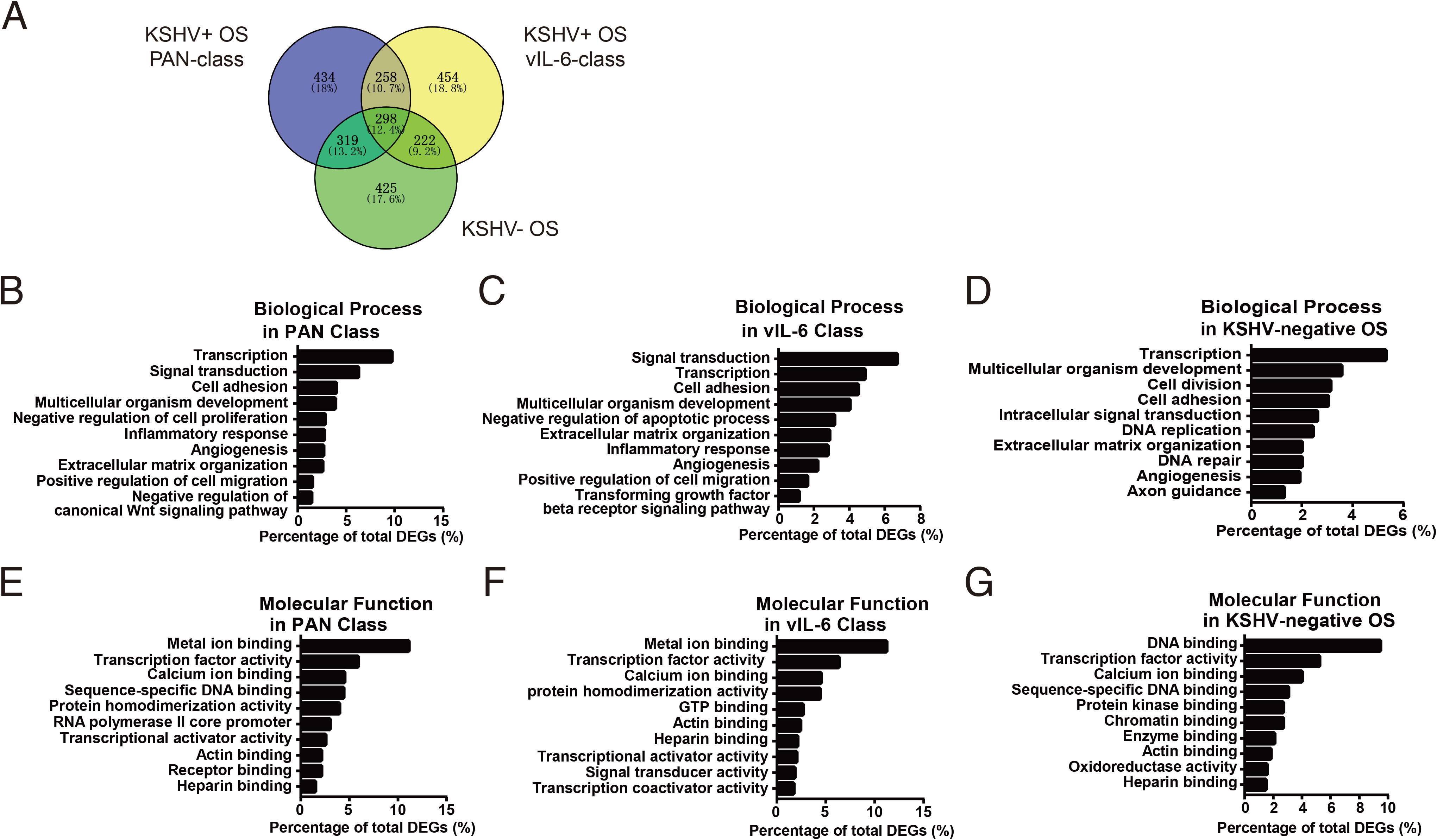
Gene ontology (GO) analysis of KSHV-positive (PAN and vIL-6 classes) and -negative osteosarcomas. (A) DEGs of KSHV-positive and -negative osteosarcomas vs. their adjacent normal tissues were analyzed and compared by Venny diagram. Numbers and percentage of DEGs unique for each of the three pairwise comparisons as well as common to two or three classes of osteosarcoma are listed. (B – D) The top 10 significantly enriched terms of biological process in DEGs of KSHV-positive osteosarcomas (PAN class, B), KSHV-positive osteosarcoma-s (vIL-6 class, C), and KSHV-negative osteosarcomas (D), vs. their adjacent normal tissues. (E – G) The top 10 significantly enriched terms of molecular function in DEGs of KSHV-positive osteosarcomas (PAN class, E), KSHV-positive osteosarcomas (vIL-6 class, F) and KSHV-negative osteosarcomas (G), vs. their adjacent normal tissues.

**Supporting Information – Fig. S3.**
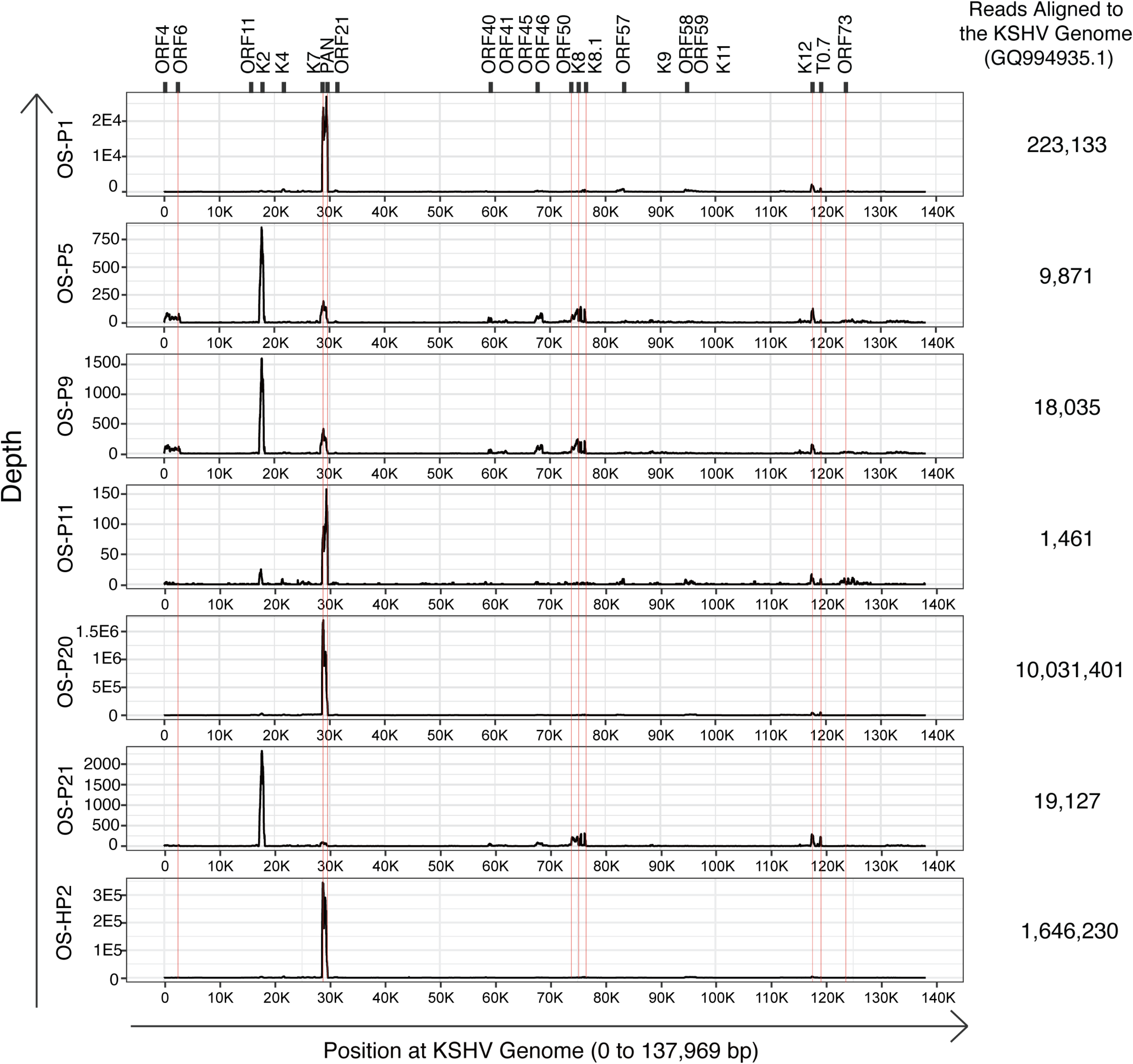
Comparison of Han osteosarcoma (OS-HP2) with six Uyghur osteosarcomas (OS-Pl, PS, P9, Pll, P20 and P21) on KSHV transcription profile. RNA-seq reads were aligned to the complete KSHV reference genome (version GQ994935.1) and visualized on a linear scale. The Y-axis represents the number of reads aligned to each nucleotide position of the KSHV genome. KSHV genome positions are indicated at the bottom of each panel. Total reads mapped to the KSHV genome ofeach clinical sample are listed on the right.

**Supporting Information – Fig. S4.**
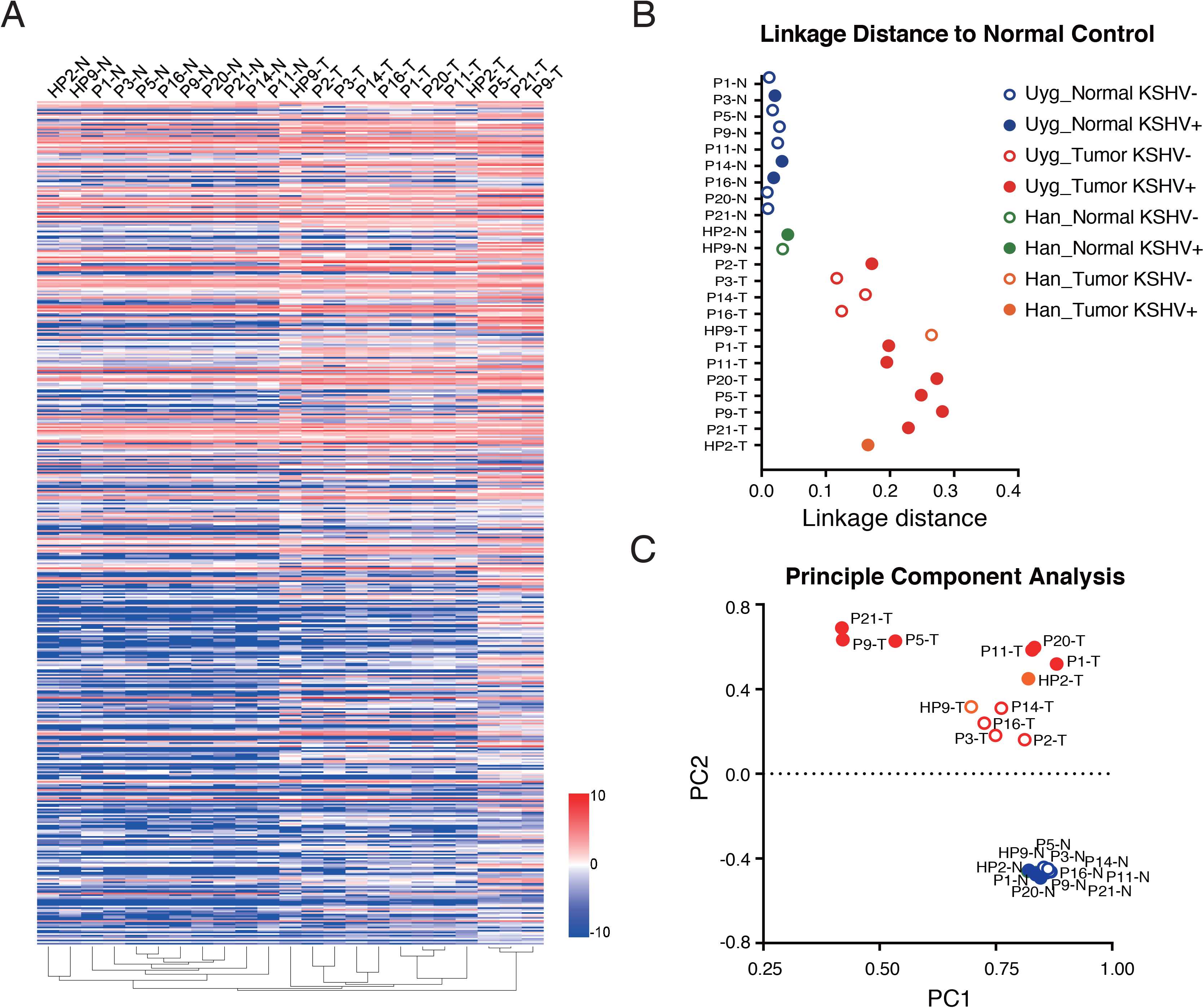
KSHV-positive and -negative osteosarcomas, regardless of Han and Uyghur patients, represent distinct subtypes of osteosarcoma on gene expression profile. (A) The RNA-seq reads of two Han (HP2 and HP9) and ten Uyghur osteosarcomas as well as their adjacent normal tissue samples were mapped to the human reference genome (version hg19/GRCh37). Unsupervised clustering of osteosarcoma samples (X-axis) and genes (Y-axis) were performed by the average linkage method. (B) Linkage distance to normal tissue was determined by the Pearson correlation coefficient. (C) The first two principal components of these data were identified and shown in multiple dimensional scaling (MDS) plot.

